# HCMV glycoprotein B, but not the pentameric complex, is a key target of antibody-mediated phagocytosis

**DOI:** 10.64898/2026.11.10.764253

**Authors:** Maura K. Farrell, Megan R. Connors, Maya Fabozzi, Kara M. Rzasa, Adelaide S. Fuller, Chelsea M. Crooks, Heather R. Martin, Philipp Kolb, Margaret E. Ackerman, Sallie R. Permar, Claire E. Otero

## Abstract

Human cytomegalovirus (HCMV) is a leading infectious cause of birth defects globally with devastating fetal health outcomes, and there is currently no HCMV vaccine available. Glycoprotein B (gB) and the pentameric complex (PC) are immunodominant HCMV envelope proteins and candidates of interest in vaccine development due to their ability to elicit strong anti-HCMV neutralization responses. Recent evidence implicates non-neutralizing antibody functions in protection from vertical HCMV transmission, but the involvement of gB and PC in these responses remains incompletely understood. We evaluated immune responses against HCMV elicited by a gB protein subunit vaccine with and without PC in a preclinical rabbit study. Despite improved neutralization and inhibition of cell-associated viral spread, the inclusion of PC did not enhance antibody-mediated phagocytosis or FcγR activation against HCMV virions or infected cells compared to gB alone. We additionally found a reduction in phagocytosis function by an HCMV hyperimmune globulin when blocked against gB, but not PC, suggesting that gB is a target of phagocytosis-mediating antibodies while PC is not. Given that antibody-mediated phagocytosis but not neutralization has been associated with protection from vertical HCMV transmission, these findings support the continued assessment of gB, but not PC, as a vaccine antigen for the prevention of vertical transmission via non-neutralizing antibodies to reduce the congenital HCMV burden worldwide.

## Introduction

Human cytomegalovirus (HCMV) is a ubiquitous beta herpesvirus, infecting roughly 83% of adults globally and nearly 1 in 3 children by the age of 5 in the U.S. [1, 2]. Congenital CMV (cCMV) impacts 1 in every 200 liveborn infants and can have detrimental results such as hearing loss, neurodevelopmental disabilities, and in extreme cases, fetal demise [3, 4]. Recently, cCMV infection was linked to increased risk of pediatric acute lymphoblastic leukemia (ALL) [5–7]. The severe impact of cCMV and high overall prevalence of HCMV makes finding an HCMV vaccine a major public health priority.

The most efficacious vaccine candidate to date is the MF59-adjuvanted glycoprotein B (gB) subunit vaccine (gB/MF59), which demonstrated 43–50% efficacy in prevention of HCMV acquisition in multiple phase II clinical trials [8–11]. This vaccine utilizes gB, a glycoprotein on the surface of the virion responsible for membrane fusion during viral entry and a dominant target of the HCMV-specific humoral response [12]. Due to the gB/MF59 vaccine’s partial success, researchers hypothesized that co-targeting gB with other glycoproteins, such as those that make up the pentameric complex (PC), would yield more complete protection [3]. PC is comprised of five viral proteins in the HCMV virion; glycoproteins H and L (gH/gL) complexed with UL128, UL130, and UL131A [3]. This complex is involved in a key mechanism of viral entry into epithelial, endothelial, and myeloid cells, but may not be essential for cell entry [3]. A recent study using a rhesus macaque model showed that PC is dispensable for cCMV infection, suggesting that PC is not necessary for CMV to cross the placenta [13]. Also, the recent phase III clinical trial for the mRNA-1647 HCMV vaccine (Moderna, NCT05085366), which contains mRNA sequences that encode for PC and gB, failed to meet the trial’s set efficacy endpoint. Despite induction of robust neutralizing antibodies, mRNA-1647 vaccination achieved 6-23% efficacy in prevention of primary infection, well below the benchmark set by the gB/MF59 vaccine [14].

In this study, we immunized rabbits, a validated model system for studying antibody Fc effector functions [15, 16], with an adjuvanted gB protein subunit vaccine similar in formulation to the gB/MF59 vaccine with and without PC added to evaluate these antigens within a consistent vaccine platform. We assessed antibody functions including neutralization, inhibition of cell-associated spread, phagocytosis, and Fc gamma receptor (FcyR) activation. We followed up on an interesting finding that inclusion of PC did not affect vaccine-elicited antibody-mediated phagocytosis activity by antigen-specific blocking of a hyperimmune globulin with high titers of HCMV-specific antibodies (Cytogam) to further evaluate gB and PC as targets for the phagocytosis-mediating antibodies in natural infection. Given that Fc effector functions have been associated with protection from vertical transmission [17, 18] an effective vaccine for prevention of cCMV will likely incorporate antigen targets of these critical responses.

## Methods

### Rabbit vaccinations and sample collection

Two-to-three-month-old New Zealand White rabbits were purchased from Charles River Laboratories (Wilmington, MA) and housed at Weill Cornell Medicine (WCM). Animals were cared for according to Guide for the Care and Use of Laboratory Animals, and all procedures were approved by the Institutional Animal Care and Use Committee (IACUC) at WCM (protocol # 2022-0017) and conducted by the Research Animal Resource Center (RARC) veterinary staff.

Vaccines were comprised of 20mcg gB (Towne strain, Sino Biological) alone or with 20mcg PC (VR1814 strain, Native Antigen Company) diluted to 250μL in sterile PBS. The protein mix was then combined 1:1 with Addavax (InvivoGen). Vaccines were administered at Weeks 0, 4, and 8 with blood draws every two weeks for 16 total weeks (**Figure 1A**). Rabbits were sedated with acepromazine 10 minutes prior to blood collection and an anesthetic cream was applied topically over the central auricular artery up to 1 hour prior to blood collection into EDTA-coated collection tubes. Vaccines were administered intramuscularly in the epaxial muscle immediately following blood collection. At Week 16, blood was collected via the central auricular artery under acepromazine sedation, followed by deep sedation then euthanasia via barbiturate overdose.

**Figure 1:**
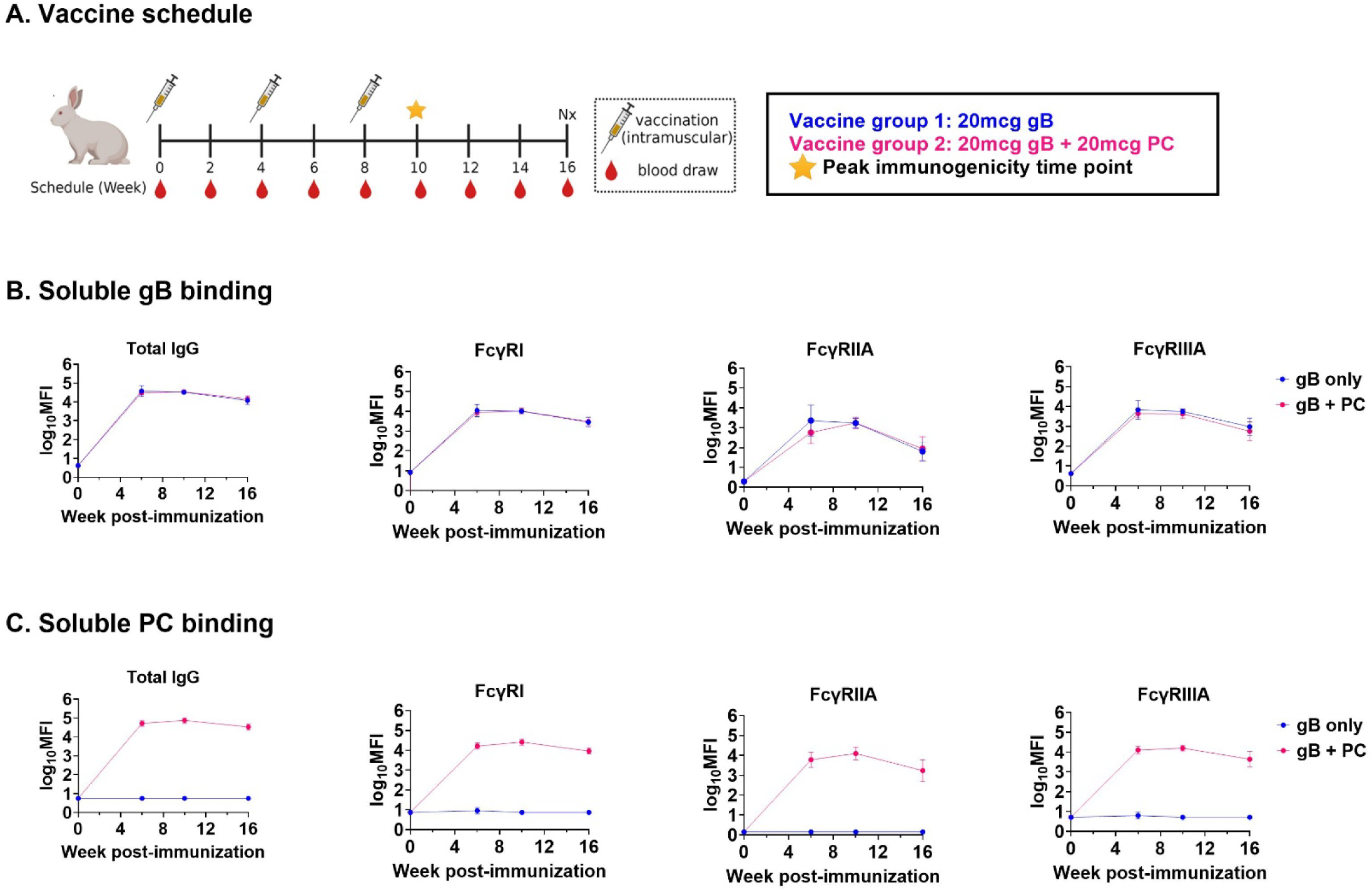
gB and PC immunization elicit antigen-specific IgG responses. (A) Rabbits were immunized via intramuscular (IM) injection with 20mcg gB (n=4) or 20mcg gB + 20 mcg PC (n=4) at weeks 0, 4, and 8. Blood was collected and processed biweekly for 16 weeks until necropsy (Nx). BAMA assessed gB and PC binding at 0, 6, 10, and 16 weeks post-immunization and FcγR binding to antigen-specific IgG. (B) Immunization with gB only and gB+PC elicited gB-specific IgG and FcγRI, FcγRIIA (H131), and FcγRIIIA (V158) binding to gB-specific IgG, reported as log_10_ of MFI minus background. (C) Immunization with gB+PC combination, but not gB-only, elicited PC-specific IgG and FcγRI, FcγRIIA (H131), and FcγRIIIA (V158) binding to PC-specific IgG, reported as log_10_MFI minus background. Error bars represent standard deviation. Schematics generated using BioRender.

Blood and tissues were processed via standard procedures. Briefly, whole blood was spun down by low-speed centrifugation to separate the plasma and cells. The plasma layer was collected and spun down again at 2000 rpm for 15 minutes to pellet any cellular contamination. Plasma aliquots were stored at −80°C until further use and heat inactivated after initial thaw by incubating at 56°C for 30 minutes.

### Binding antibody multiplex assay (BAMA)

BAMA was performed as described previously [10, 18–21]. MagPlex magnetic carboxylated fluorescent beads (Luminex) with different fluorescent signatures (bead regions) were covalently coupled to individual antigens. Coupled beads were incubated with rabbit plasma (1:5000) or Cytogam (0.005-111.1 mcg/mL). Antigen-specific antibody binding was detected with recombinant biotinylated human Fc*γ*RI, FcγRIIA (H131), or FcγRIIIA (V158) (1mcg/mL, produced by Duke Protein Production Facility and biotinylated in-house) incubated with phycoerythrin (PE)-conjugated streptavidin (BD Biosciences). Antigen-specific antibody binding for total IgG was detected with PE-conjugated goat anti-rabbit IgG (2mcg/mL, Southern Biotech) for rabbit samples or PE-conjugated goat anti-human IgG (2mcg/mL, Southern Biotech). Total IgG and FcγR binding against purified gB (Towne strain, Sino Biologicals) and PC (AD169 strain, received from Jason McLellan) were measured on the xMAP Intelliflex system (Luminex). Antigen-specific antibody binding for total IgG with gH/gL (TB40e strain, produced in-house) and gB AD-4, AD-5, AD-4+5, and AD-1 (Merlin strain, produced in-house, **Table S1**) was measured on the Bio-Plex 200 system (Bio-Rad) with plasma diluted 1:2000. A diluent-only sample was included on each plate to determine bead-specific background binding and reported median fluorescence intensities (MFIs) were background subtracted. A serial dilution of HCMV hyperimmune globulin (Cytogam, CSL Behring) and a point dilution of palivizumab (Synagis) were included on each plate as positive and negative controls, respectively, and as an assessment of inter-plate variability. Antigen-specific antibody binding for HCMV hyperimmune globulin and palivizumab was detected using PE-conjugated goat anti-human antibody specific for IgG (2 mcg/mL, Southern Biotech). A seropositive rabbit plasma sample and a point dilution of a rabbit IgG isotype control were also included as additional positive and negative controls. All samples were run in duplicate, and results are reported as the average of both replicates. Any sample with a blank bead MFI >1000, an antigen with a CV over 30%, or bead count below 50 was repeated one time. A positivity threshold was calculated for each antigen as the average + 2 SD of the week 0 plasma samples. Samples falling below the positivity threshold were set to half of the positivity threshold.

### IL-2 reporter cell FcγR activation

FcγR activation was measured using human-FcγR-expressing BW5147 IL-2 reporter cells as previously described [22] targeting recombinant proteins (gB, Sino Biologicals, 4mcg/mL; PC, Native Antigen Company, 2mcg/mL), AD169r virions (500 pfu/well), or AD169r-infected HFF-1 cells (1MOI, 48h). Following 1-2h incubation with plasma/antibody samples and subsequent overnight incubation with the reporter cells (20,000 cells/well), IL-2 in the supernatant was quantified using a sandwich ELISA for mouse IL-2 as a surrogate marker of activation of the engineered FcγR expressed on each reporter cell line. A serial dilution of HCMV hyperimmune globulin (Cytogam, CSL Behring) and a point dilution of a rabbit IgG isotype control were included on each plate as positive and negative controls, respectively, and as an assessment of inter-plate variability. A mouse IL-2 positive control was included on each ELISA plate to ensure consistency of the ELISA readout between plates. Each sample was run with two biological replicates, each with two technical ELISA replicates. A positivity threshold was calculated for each antigen as the average + 2 SD of the week 0 plasma samples. Data passed quality control checks if samples above the positivity cutoff had CVs below 30%.

### ELISA

ELISA was performed as described in Karthigeyan et al. with some modifications [20]. 384-well ELISA plates were coated with gB antigenic domain 2 (AD-2) site 1 (AD169 Strain; HRANETIYNTTLKYG) or gB antigenic domain 6 (AD-6, Towne strain, CMIALDIDPLENTDFRVLELYSQKELRSSNVFDLEEIMREFNSYKQRVKYV) (1mcg/well, Genscript) overnight at 4°C, then blocked in assay diluent (1× PBS 4% whey, 15% normal goat serum, and 0.5% Tween 20) for 1 hour at room temperature. Rabbit plasma samples (1:50, followed by 8-point 6-fold serial dilution) were added for 1 hour at room temperature to allow for antigen binding, followed by a 1-hour incubation with horse radish peroxidase-conjugated goat anti-rabbit IgG secondary (1:5000, Southern Biotech) to detect binding. Plates were developed using SureBlue Substrate (SeraCare) followed by addition of equal volume of Stop Solution (SeraCare), and optical density was measured using a BioTek Synergy plate reader. All samples were run in duplicate and repeated if positive readings demonstrated greater than 20% CV between replicates. Results were reported as log AUC over the plasma dilution series.

### Neutralization

As described previously, retinal pigment epithelial cells (ARPE-19, ATCC) and human foreskin fibroblasts (HFF-1, ATCC) were seeded 6,000 cells per well in 384-well plates (Corning, 3764) and incubated at 37°C overnight [20]. Samples were heat inactivated and serially diluted in cell growth media, with and without standard rabbit complement (Cedar Lane, CL310) diluted 8-fold in cell growth media [10, 23]. AD169r (multiplicity of infection [MOI] =1 for epithelial cells and MOI=2 for fibroblast cells) was incubated with plasma for 1 hour at 37°C in 96 well plates. The virus-plasma dilutions were added to the 384-well plates with confluent HFFs for 24 hours and ARPEs for 48 hours. Plates were fixed with 10% formalin and stained with mouse anti-IE-1 (Sigma Aldrich), followed by anti-mouse IgG-AF488 (abcam) to detect infected cells and 4’,6-diamidino-2-phenylindole (DAPI) for nuclei. In addition to no virus and virus only cell controls, a serial dilution of HCMV hyperimmune globulin (Cytogam, CSL Behring) and a point dilution of palivizumab (Synagis) were included on each plate as positive and negative controls, respectively, and as an assessment of inter-plate variability. Plates were imaged and analyzed on the ImageXpress Pico Automated Cell Imaging System (Molecular Devices). The 50% inhibitory doses (ID_50_) were calculated on GraphPad Prism version 10.4.1 (GraphPad Software, Inc, LaJolla, CA) using nonlinear regression (Sigmoidal, 4PL) with average virus control set as the top constraint and 0 as the bottom.

### Cell-To-Cell Associated Spread

Similar to previous studies, prevention of cell-associated viral spread was assessed by infecting confluent ARPE-19 cells with Ts15nr (MOI=0.1) in 384 well plates [20]. After 48 hours, cells were washed with PBS fortified with 1% fetal bovine serum (FBS). Serum from vaccinated rabbits was diluted 1:40 in cell growth medium, added to cells in duplicate, and incubated at 37°C for 12 days. In addition to no virus and virus only cell controls, a serial dilution of HCMV hyperimmune globulin (Cytogam, CSL Behring) and a point dilution of palivizumab (Synagis) were included on each plate as positive and negative controls, respectively, and as an assessment of inter-plate variability. Plates were stained and imaged as in the neutralization assays described above and results reported as the percent reduction in infection by each sample relative to the virus-only control wells.

### Antibody-mediated phagocytosis

Virion phagocytosis activity was assessed as described previously [18]. HCMV strains AD169r (125 PFU/well), Towne (125 PFU/well), or TB40e (250 PFU/well) were conjugated to (10 mcg) AF647 (Invitrogen) and quenched with Tris-HCl (pH 8.0) after two hours of constant agitation at room temperature. In 96-well plates, rabbit plasma (1:30) was incubated with AF647-conjugated HCMV at 37°C for two hours. THP-1 monocytes (ATCC) were then added at 50,000 cells per well before spinning plates at 1200g for 1 hour at 4°C. Then, plates were incubated for 1 hour at 37°C. Cells were stained with Aqua live/dead stain (ThermoFisher), washed, fixed with 10% formalin (Fisher Brand), and resuspended in PBS before being acquired on a BD LSR Fortessa. Phagocytosis was quantified by gating for the percentage of single, live, AF647+ THP monocytes using FlowJo software (**Figure S2**). In addition to no virus and virus only cell controls, a serial dilution of HCMV hyperimmune globulin (Cytogam, CSL Behring) and a point dilution of palivizumab (Synagis) were included on each plate as positive and negative controls, respectively, and as an assessment of inter-plate variability.

Phagocytosis targeting recombinant antigen coated beads was assayed as previously described [24] with minor modifications. Recombinant HCMV gB (Sino Biological) and CMV gH pentamer complex (Native Antigen Company) were biotinylated (Thermo) before conjugation to FluoSpheres NeutrAvidin microspheres (Invitrogen F8776) for 20 minutes with end over end mixing at room temperature. Post conjugation they were spun down and washed twice with PBS-BSA. Conjugated beads were stored for a few days at 4°C prior to use. Cytogam was assayed at concentrations ranging from 15.625 to 2,500 mcg/mL and incubated with the FluroSpheres for 30 mins at room temperature before 2×10^4^ cells per well were added, yielding a final bead to cell ratio of 40:1 and final volume of 100 µL. The plates were then incubated at 37°C for 5 hours before being fixed with 4% paraformaldehyde (Electron Microscopy Sciences). Once fixed, they were washed and resuspended in PBS (Corning) before data acquisition on an Agilent NovoCyte flow cytometer (NovoExpress Version 1.6.2). Data was analyzed with FlowJo (Version 10.9.0). Data was plotted as phagocytosis score, which was calculated by (% bead positive cells x geometric mean MFI).

To assess antigen specificity, a virion phagocytosis blocking assay was performed. In 96-well plates, Cytogam (50mcg/mL) was incubated with a serial dilution of soluble gB or PC (7.81-250mcg/mL) or CTLA4, a negative control blocking protein. The rest of the assay was performed as previously described.

### Statistical analysis

Data was analyzed through GraphPad Prism version 10.4.1 (GraphPad Software, Inc, LaJolla, CA). Comparisons of antibody kinetics in vaccine groups was assessed using a two-way repeated measures ANOVA on vaccine group and time. Sidak’s test was used to compare vaccine groups at each time point when a significant vaccine*time interaction term (p<0.05) was observed in the overall ANOVA and reported as significant for adjusted p-values below 0.05. For assays focused on peak immunogenicity at Week 10, Mann-Whitney tests were used to compare each response between vaccine groups.

## Results

### gB and PC immunization elicit antigen-specific IgG responses

Rabbits were immunized in a three-dose series with 20mcg gB with or without 20mcg PC (n=4 each), adjuvanted with Addavax (**Figure 1A**). Plasma IgG binding responses and human FcγR binding to antigen-specific IgG were assessed at weeks 0, 6, 10, and 16 via BAMA. gB-only and gB+PC immunization elicited strong IgG binding responses to gB (**Figure 1B**), and gB+PC immunization elicited strong IgG binding responses to PC (**Figure 1C**). We found no difference in FcyRI, FcyRIIA (H131), or FcyRIIIA (V158) binding to gB-specific IgG between gB-only and gB+PC vaccine groups (**Figure 1B**). The gB+PC vaccine group, but not the gB-only group, showed FcyRI, FcyRIIA, and FcyRIIIA binding to PC-specific IgG (**Figure 1C**). Antigen-specific FcyR activation was assessed using an IL-2 reporter cell assay targeting recombinant gB and PC [22], which confirmed that gB and gB+PC immunization elicited gB-specific IgG capable of activating FcyRI, FcyRIIA, and FcyRIIIA, and identified faster waning of FcγRIIIA signal induction activity (repeated measures two-way ANOVA vaccine*time interaction p=0.008, Sidak’s p[Week 16]=0.04) but no other significant differences between vaccine groups. Only the combination vaccine elicited PC-specific IgG capable of activating these FcyRs (**Figure 2**).

**Figure 2:**
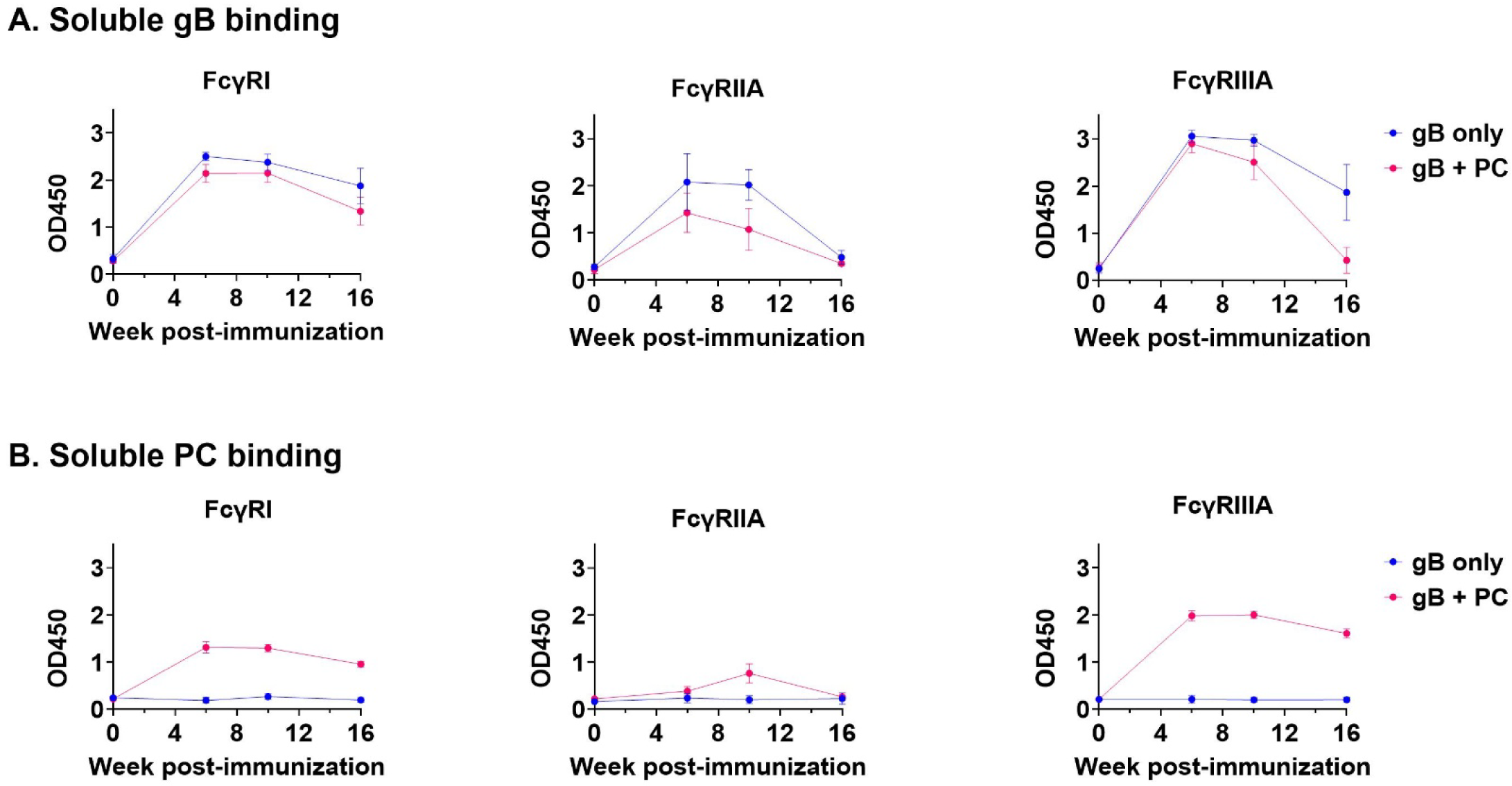
FcγR activation confirms the presence of vaccine elicited gB- and PC-specific IgG. An antigen-specific FcγR activation assay was used to assess vaccine-elicited gB- and PC-specific IgG measured by IL-2 release of reporter cells using ELISA and reported as OD450. A repeated measures two-way ANOVA was used to compare vaccine groups over time, with Sidak’s pairwise comparisons following a significant (p < 0.05) vaccine*time interaction term. (A) gB only and gB+PC groups both produced gB-specific IgG capable of activating FcγRI, FcγRIIA (H131), or FcγRIIIA (V158). Only FcγRIIIA demonstrated a significant interaction between vaccine group and time (p=0.008), which was driven by the difference at Week 16 (p=0.04). (B) gB+PC groups, but not gB only groups, produced gB-specific IgG capable of activating FcγRI (p<0.0001), FcγRIIA (p=0.0002), or FcγRIIIA (p<0.0001). For FcγRI and FcγRIIIA, all time points after Week 0 demonstrated a significant difference between groups (p≤0.0001). Error bars represent standard deviation. FcγRIIA only demonstrated significantly different activation at Week 10 (p=0.03).

We additionally measured IgG binding responses against components of each vaccine antigen at Week 10. Similar to observations of total IgG binding to the whole, soluble PC and gB antigens, we observed IgG responses against gH/gL, the glycoprotein complex forming the base of the PC, only in the gB+PC vaccine group (**Figure 3A**). We additionally observed similar IgG binding responses against each AD of gB (excluding the cytosolic AD-3) in both vaccine groups. Notably, IgG binding was detected well above background against AD-1, AD-4, AD-5, and AD-4+5 (**Figure 3B-E**) but only a subset of animals developed IgG responses exceeding background against AD-2 site 1 and AD-6 (**Figure 3F-G**).

**Figure 3:**
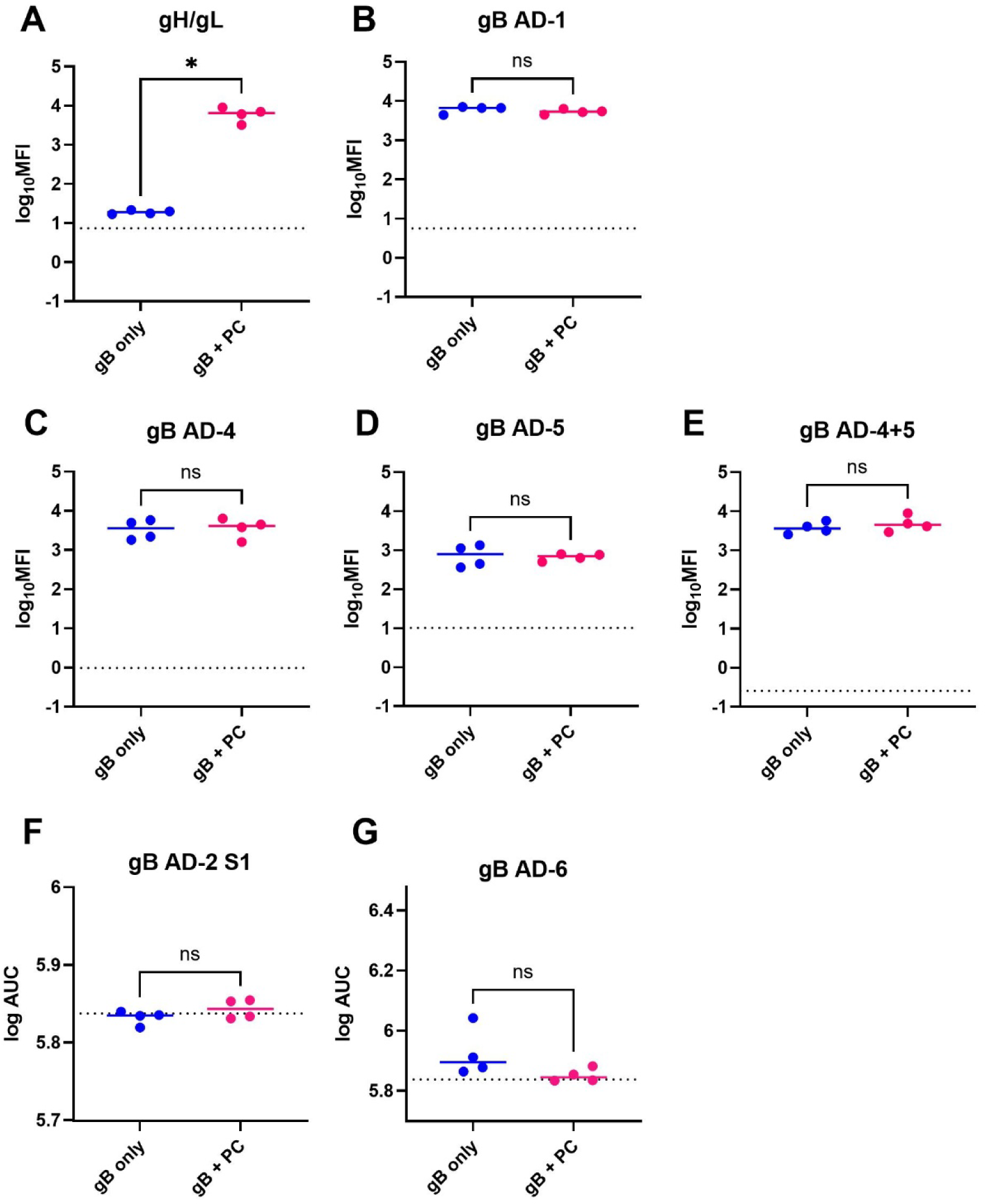
IgG responses against components of each vaccine antigen. We used BAMA to assess IgG binding to (A) gH/gL, a major component of PC, and (B-E) antigenic domains (ADs) of gB, reported as log_10_ of MFI minus background. Due to the size of the peptides and incompatibility with coupling to BAMA beads, we utilized ELISAs to assess IgG binding responses against (F) gB AD-2 site 1 and (G) AD-6, represented as the log AUC over the plasma dilution series. Dashed lines represent week 0 baseline. Mann-Whitney tests were used to statistically compare groups.

### The addition of PC to a gB vaccine enhances neutralizing antibody responses but not Fc mediated effector responses against HCMV

Neutralization assays were performed with AD169r HCMV strain on fibroblast and epithelial cells to test the functional responses of the vaccines 10 weeks post vaccination. Overall, plasma samples from rabbits vaccinated with PC demonstrated significantly higher HCMV neutralization on epithelial cells than rabbits vaccinated with only gB in the absence or presence of rabbit complement (**Figure 4A-B**, Mann-Whitney p=0.03 and p=0.03, respectively). The gB+PC vaccine also elicited higher neutralizing-antibody responses against AD169r on fibroblasts than the gB vaccine in the presence of rabbit complement, but the magnitude of this difference was notably lower than on epithelial cells (**Figure 4C**, Mann-Whitney p=0.03). A cell-associated viral spread assay compared the capacity of the gB only and gB+PC vaccines to induce antibodies that block cellular dissemination of the Ts15nr strain of HCMV on epithelial cells at Week 10 (**Figure 4D**). The gB+PC vaccine immunized rabbits elicited responses that prevented viral spread significantly better than the gB only group (Mann-Whitney p=0.03).

**Figure 4:**
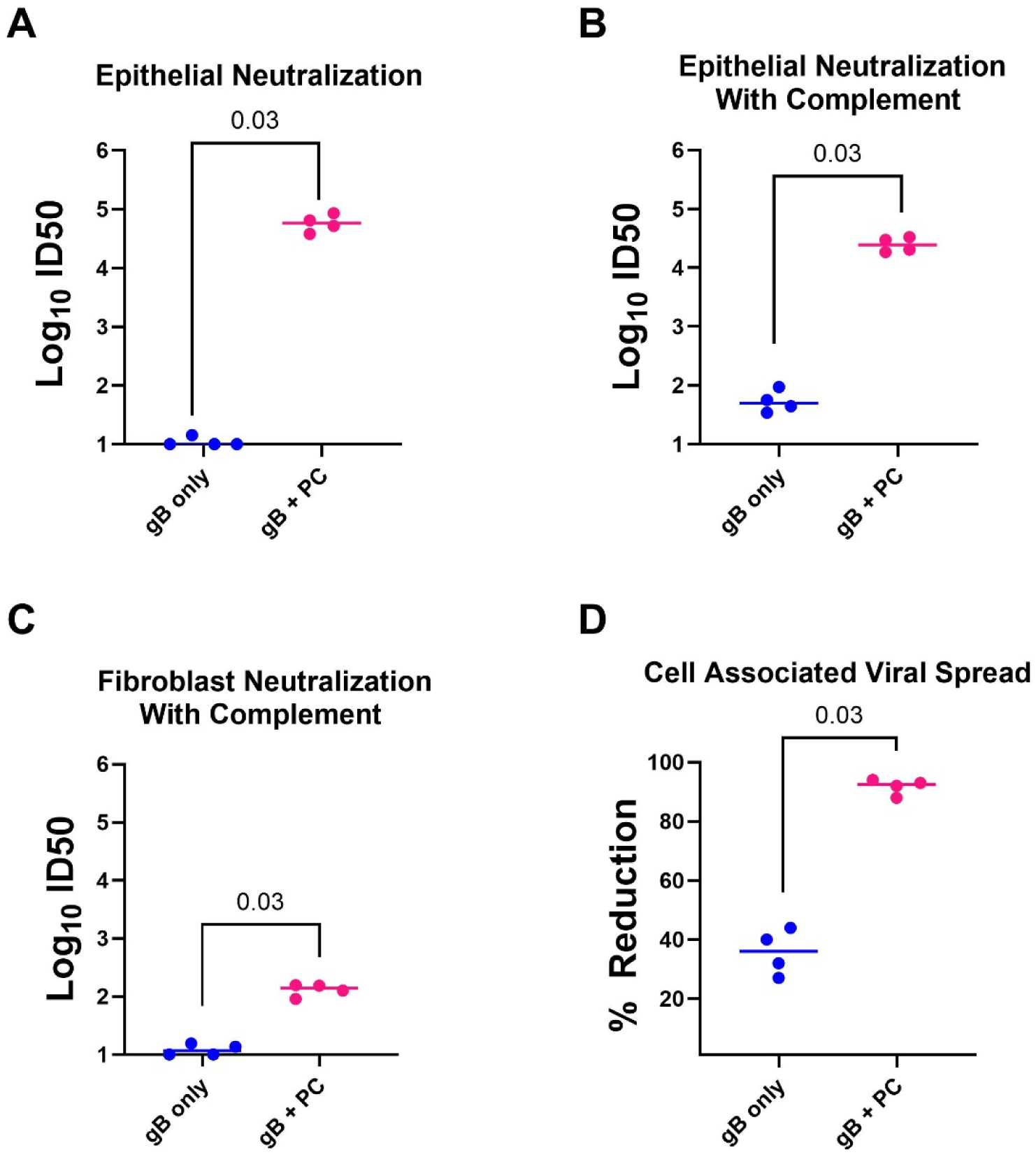
The addition of PC elicited greater levels of neutralizing antibodies against HCMV than gB on its own. The neutralization assay assed neutralizing antibodies against AD169r elicited in rabbit plasma 10 weeks post-immunization. (A-B) The gB+PC group elicited antibodies that neutralized AD169r on epithelial cells, with or without complement, significantly better than the gB only group (Mann-Whitney p=0.03 and p=0.03, respectively). (C) The gB+PC group elicited antibodies that neutralized AD169r on fibroblast cells in the presence of rabbit complement significantly better than the gB only group (Mann-Whitney p=0.03). (D) The inhibition of cell-associated virus spread assay assessed reduction of viral spread of the Ts15nr strain on epithelial cells by immunized rabbit plasma diluted 1:40. Cell-associated spread of Ts15nr on epithelial cells was reduced by responses elicited by vaccination with gB+PC significantly more than gB only (Mann-Whitney p=0.03).

Antibody-mediated phagocytosis activity of rabbit plasma was assessed at 10 weeks post-immunization with gB and gB+PC. Phagocytosis activity against AD169r and Towne was above week 0 baseline and not significantly different between groups immunized with gB versus gB+PC (**Figure 5A**). FcyR activation was assessed using an IL-2 reporter cell assay[22]. Activation of FcyRI, FcyRIIA, and FcyRIIIA by Week 10 plasma antibodies targeting AD169r virions were not above baseline and not significantly different between groups immunized with gB versus gB+PC (**Figure 5B**). This was also the case when we assessed week 10 plasma antibodies against AD169r-infected cells (**Figure 5C**).

**Figure 5:**
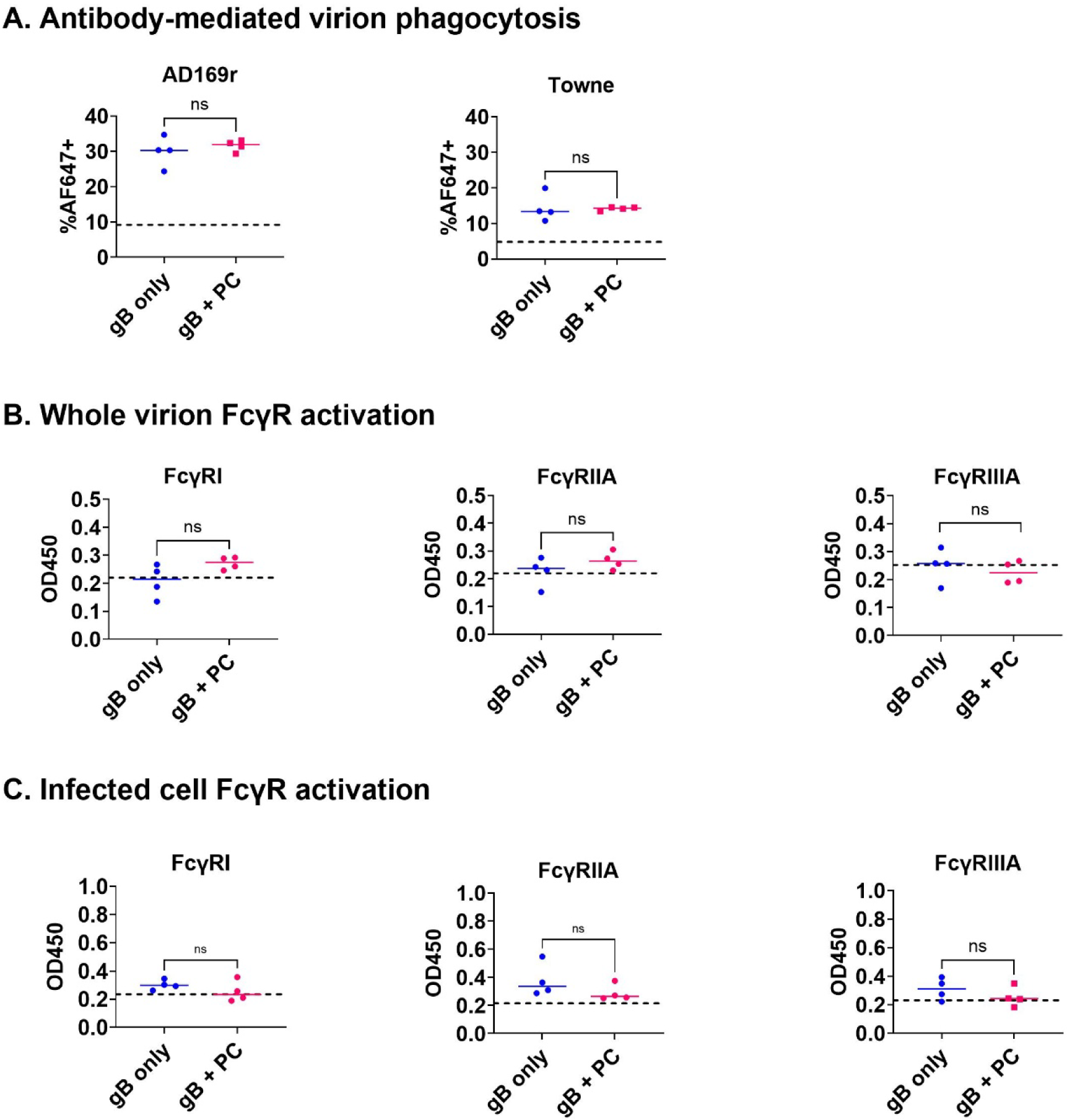
PC does not enhance antibody mediated phagocytosis or FcγR activation against HCMV. (A) Antibody-mediated phagocytosis activity against AF647-conjugated HCMV mediated by plasma from rabbits 10 weeks post-immunization with 20mcg gB or 20mcg gB + 20mcg PC, reported as %AF647+ THP monocytes. There was no difference in antibody-mediated phagocytosis activity against AD169r or Towne between gB-only or gB+PC groups. FcγR activation of reporter cells co-cultured with (B) AD169r virions or (C) AD169r-infected cells and plasma from rabbits 10 weeks post-immunization measured by IL-2 release using ELISA and reported as OD450. There was no difference in activation of FcγRI, FcγRIIA (H131), or FcγRIIIA (V158) between gB-only or gB+PC groups. Dashed lines represent week 0 baseline. Mann-Whitney tests were used to statistically compare groups.

### gB, but not PC, effectively blocks virion phagocytosis of an anti-CMV hyperimmune globulin

To further explore our finding that the inclusion of PC in a gB protein subunit vaccine did not enhance virion phagocytosis responses, we evaluated the involvement of each antigen in the antibody-mediated phagocytosis response to natural infection using Cytogam, a hyperimmune globulin isolated from HCMV-seropositive individuals, which binds gB and PC in a dose-dependent manner (**Figure 6A**).

**Figure 6:**
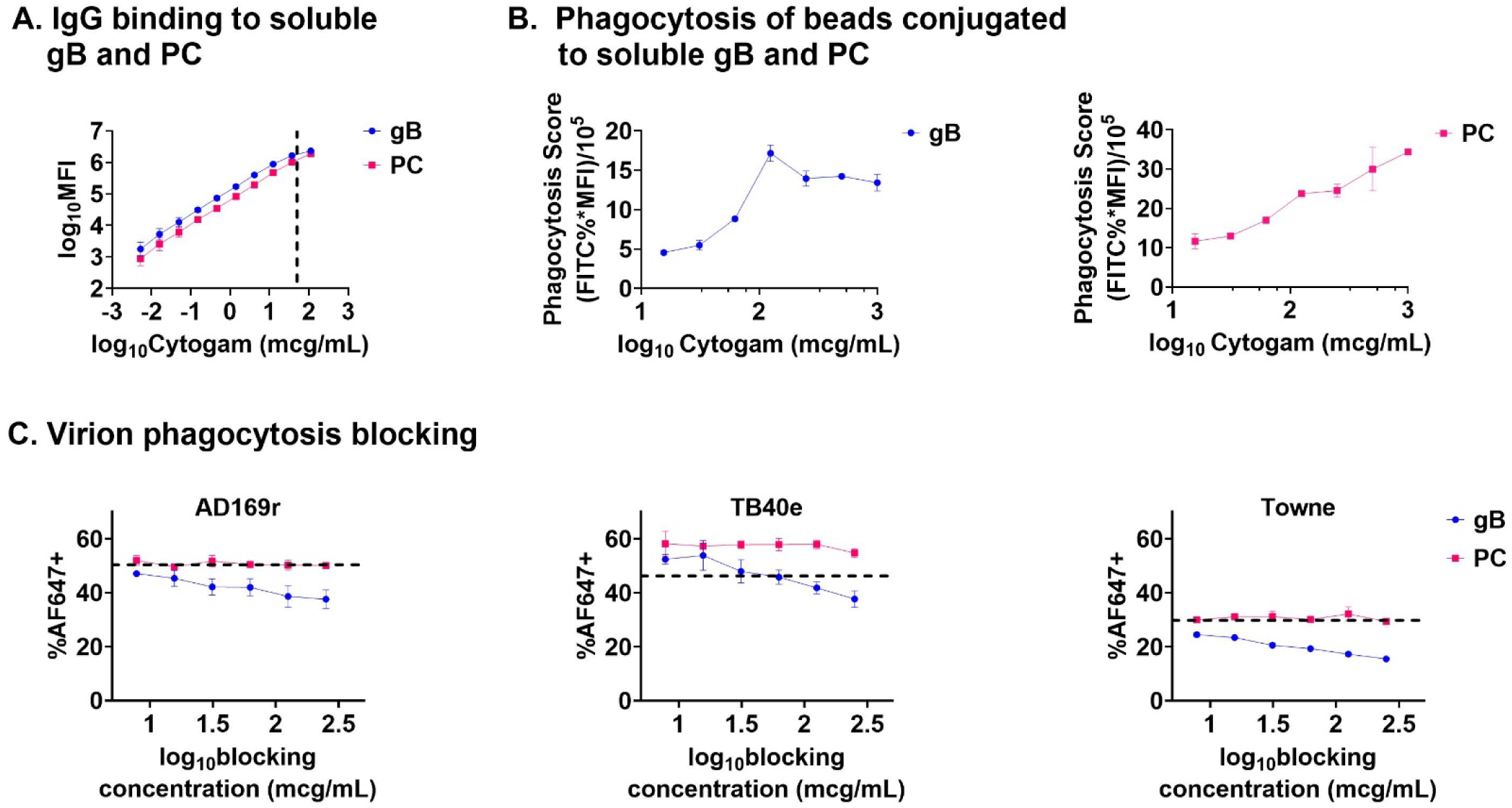
Soluble gB, but not PC, blocks phagocytosis activation by an anti-CMV hyperimmune globulin. (A) Cytogam (0.05-3000 mcg/mL) binds gB and PC in a dose-dependent manner, measured via BAMA. Dashed line indicates Cytogam at 50mcg/mL as used in phagocytosis blocking assays. (B) Cytogam (15.625-2500 mcg/mL) mediates phagocytosis of gB- and PC-conjugated beads, reported using a phagocytosis score (FITC%*MFI)/10⁵). (C) Virion phagocytosis activity of Cytogam was measured in the presence of soluble gB and PC. Phagocytosis activity against AD169r, TB40e, and Towne decreased in a dose-dependent manner in the presence of soluble gB, but not PC. Error bars represent standard deviation. Dashed lines indicate phagocytosis activity of unblocked Cytogam at 50mcg/mL.

To assess the targeting of gB and PC in anti-HCMV phagocytosis responses, we used bead-based and competitive blocking approaches. We found that Cytogam mediates phagocytosis of both gB- and PC-conjugated beads in a dose-dependent manner (**Figure 6B**). However, in a competitive blocking assay, phagocytosis activity of Cytogam against AD169r, TB40e, and Towne virions decreased in a dose-dependent manner only in the presence of gB (**Figure 6C**). Virion phagocytosis activity showed no reduction or dose effect in the presence of PC (**Figure 6C**).

## Discussion

In this report, we show that gB, but not PC, is a key target of the antibody-mediated phagocytosis response against HCMV in immunized rabbits and in pooled human serum immunoglobulin from CMV seropositive donors, supporting recent studies that call into question the utility of PC as an HCMV vaccine antigen [13, 14]. In our preclinical rabbit vaccine study, IgG binding responses to gB were similar between both groups, and only the group immunized with gB+PC developed PC-specific IgG as expected. Both gB- and PC-specific IgG engaged with human FcγRI, FcγRIIA, and FcγRIIIA, suggesting the ability to mediate Fc effector functions.

However, FcγR activation against HCMV virions and infected cells was low in both groups. Additionally, virion phagocytosis responses were measurable and similar in both groups, suggesting that inclusion of the additional glycoprotein antigen targets in PC did not improve elicited antibody-mediated virion phagocytosis. This finding was further supported by competitive blocking experiments using pooled IgG from HCMV-seropositive individuals, which found a reduction in phagocytosis function when blocked with gB but not PC.

Neutralization is characterized as an in vitro function of antibody mediated by binding viral antigens by their antibody binding (Fab) regions and can be mediated in the absence of the (Fc) region [25, 26]. In alignment with a previous study comparing the immunogenicity of gB/MF59 vaccine to the gB+PC-containing mRNA-1647 vaccine, gB+PC mRNA vaccination elicited greater titers of neutralizing antibodies than the gB-only protein vaccine [21]. Accordingly, as we have previously observed with gB immunizations in rabbits [10], gB alone elicited weak neutralizing antibody responses, which were markedly improved with the addition of PC. It is likely that the gB+PC vaccine elicited better neutralizing antibody responses on epithelial cells because HCMV uses PC to enter non-fibroblast cells [27, 28]. The marginal improvement in neutralization on fibroblasts may have been a result of antibodies targeting components of the pentamer, such as gH and gL, that are likely involved in entry into multiple cell types [28]. One drawback of this study is that we cannot confirm if these neutralizing antibodies are elicited only against PC or if there may have been a synergistic effect of the two antigens working in tandem. While the significant neutralizing responses favor inclusion of PC in a vaccine, multiple vaccine trials have shown that neutralization alone does not determine a HCMV vaccine’s success [10, 18].

While there is a lack of evidence surrounding neutralization as a primary correlate of protection against HCMV acquisition or congenital transmission, there is evidence that the quality of the antibody responses are critical to HCMV vaccine success [29–32]. Thus, HCMV vaccine studies have begun looking at a vaccine’s ability to induce other types of antibody functions, including inhibition of cell-to-cell virus spread as a more clinically relevant predictor [33]. Cell-to-cell spread is especially of interest because HCMV often disseminates via direct spread between neighboring cells more often than the release of cell-free virus [28, 33–35]. The mechanisms of HCMV cell-to-cell spread have yet to be fully understood, so in this study we use an operational definition of cell-associated spread [33]. We compared the gB only group and the gB+PC group’s capacity to reduce cell-associated spread of the highly cell-associated Ts15nr HCMV strain on epithelial cells. A prior study attributed the partial success of the gB/MF59 vaccine in transplant recipients to inducing responses against the antigenic domain (AD)-6 epitope, which have been associated with inhibition of cell-to-cell spread of HCMV [34].

However, responses against AD-6 were low in both groups, with the gB only vaccine group demonstrating a greater frequency of detectable anti-AD-6 IgG. Since the group vaccinated with gB+PC reduced viral spread in epithelial cells significantly better than the gB only group, PC is likely utilized in a key mechanism of viral entry into epithelial cells, both for cell-free virus and cell-to-cell spread [36].

In a cohort of HCMV transmitting and non-transmitting pregnancies, elevated antibody-mediated virion phagocytosis responses were associated with a decreased risk of vertical transmission [18], suggesting that phagocytosis plays a protective role against congenital HCMV infection. Furthermore, while both were below the level of activity observed in HCMV-seropositive individuals, antibody-mediated phagocytosis was higher in HCMV-naïve gB/MF59 vaccinees versus mRNA-1647 vaccinees, which may provide some clue as to the relative success of these two vaccines [21]. Given these clinical findings, developing an HCMV vaccine that elicits antibodies capable of mediating phagocytosis is of strong interest. Cytogam mediated phagocytosis of gB- and PC-conjugated beads, suggesting that natural infection elicits antibodies capable of mediating phagocytosis against both antigens. However, the addition of PC did not further enhance gB vaccine-elicited virion phagocytosis responses in our vaccine study, and soluble gB, but not PC, blocked phagocytosis mediated by Cytogam. These results suggest that, despite the presence of phagocytosis-mediating antibodies, other factors such as steric hindrance and complexity of glycoproteins on the viral envelope may disqualify PC as a physiologically relevant phagocytosis target. One potential limitation is that we only measured phagocytosis of fluorophore-tagged HCMV virions and not infected cells, which is another relevant immune response [37]. Despite this limitation, our study indicates that gB is a target for antibody-mediated phagocytosis in both vaccine-elicited and natural immunity, supporting the continued exploration of gB for a vaccine that enhances Fc effector responses.

In the same cohort of HCMV transmitting and non-transmitting pregnancies, elevated antibody-dependent cellular cytotoxicity (ADCC)-mediating antibodies and engagement of FcyRI, FcyRIIA and FcyRIIIA were also associated with a decreased risk of vertical transmission [17, 18]. FcyRI, FcyRIIA, and FcyRIIIA are known to mediate phagocytosis and ADCC, among other important immune functions [38]. Given this, we were interested in exploring activation of these receptors post-immunization with gB and gB+PC. Consistent with phagocytosis results, we show that the addition of PC to a gB vaccine does not enhance FcyR activation.

However, FcyR activation against HCMV virions and infected cells was not detected above baseline in either immunization group. This low level of FcyR signaling may be explained by the expression of viral Fc gamma receptors (vFcyRs) by virions and infected cells, which could bind the Fc region of gB- and PC-specific IgG, preventing them from activating FcyRs on the reporter cells [22, 39]. Another potential limitation is the use of reporter cells that express individual FcyRs, which, while providing a valuable method for assessing FcyR activation, may fail to reflect the complexity of immune cell surface FcyR interactions with anti-HCMV antibodies [22]. Lastly, surface proteins on the mouse reporter cell line, which would not otherwise be expressed on human leukocytes, may contribute to steric hindrance, making anti-HCMV antibodies less accessible for FcyR binding. Future studies should assess other immune cell-mediated Fc-mediated functions, such as ADCC responses, to determine if PC enhances gB vaccine-elicited cytotoxicity.

Overall, we did not observe that the addition of PC to a gB vaccine further enhanced non-neutralizing antibody responses against HCMV. Given the implication of Fc effector functions but not neutralization in protection from HCMV-associated disease, these findings support the continued inclusion of gB, but not PC, as a candidate vaccine antigen. While gB and PC are two of the most well-studied envelope proteins, other proteins, such as the newly defined GATE complex (gH/UL116/UL141), should be further assessed as potential vaccine antigens due to their demonstrated role as targets of ADCC-mediating antibodies [40]. By continuing to explore non-neutralizing antibody responses in vaccine development, we aim to develop a safe and effective vaccine that prevents vertical transmission and reduces the global burden of congenital HCMV.

### Conclusion

Consistent with prior studies of gB immunizations [15], we report relatively low levels of HCMV neutralizing antibodies and robust antibody-mediated virion phagocytosis responses elicited by gB alone. Inclusion of PC enhanced titers of neutralizing antibodies as well as those inhibiting cell-associated spread of HCMV but did not affect the phagocytosis response, despite evidence that both gB- and PC-specific antibodies can bind and activate human FcγRs. We followed up on this finding to evaluate gB and PC as targets of antibody-mediated phagocytosis in natural infection, utilizing Cytogam as a model antibody source. Consistent with our findings in the rabbit model, we observed strong binding responses, engagement of human FcγRs, and phagocytosis of antigen coated beads. However, utilizing an antigen blocking approach, we observed a strong dose-dependent decrease in virion phagocytosis when Cytogam was blocked against gB but no effect when blocked against PC. Overall, our results suggest that gB, but not PC, is a key target for phagocytosis-mediating antibodies.

Given that antibody-mediated phagocytosis, but not neutralization, was found to be associated with protection from vertical HCMV transmission, this is likely a key response to elicit by vaccination and, as such, PC may not be a strong candidate antigen for further vaccine development for prevention of congenital HCMV.

## Acknowledgements

We would first like to acknowledge the rabbits that were used in this study and the excellent care that they received from RARC veterinary and husbandry staff. Thanks to the Laboratory of Comparative Pathology for conducting tissue collection at necropsy. We additionally acknowledge the Flow Cytometry Core at WCM for their support of this project.

## Data Availability Statement

All data used in the present study are available upon request to the authors.

