## Supplemental Materials for "HCMV glycoprotein B, but not the pentameric complex, is a key target of antibody-mediated phagocytosis"

| TABLE S1: gB antigenic domain sequence information for BAMA antigens | |
|---|---|
| **AD** | **Residues*** |
| gB AD-1 | 549–645 |
| gB AD-4 | 112–132 and 343–438, Ile-Ala-Gly-Ser-Gly flexible linker |
| gB AD-4+5 | 112–438 |
| gB AD-5 | 133–343 |
| *All from HCMV gB Merlin strain (UniProtKB: F5HB53) | |

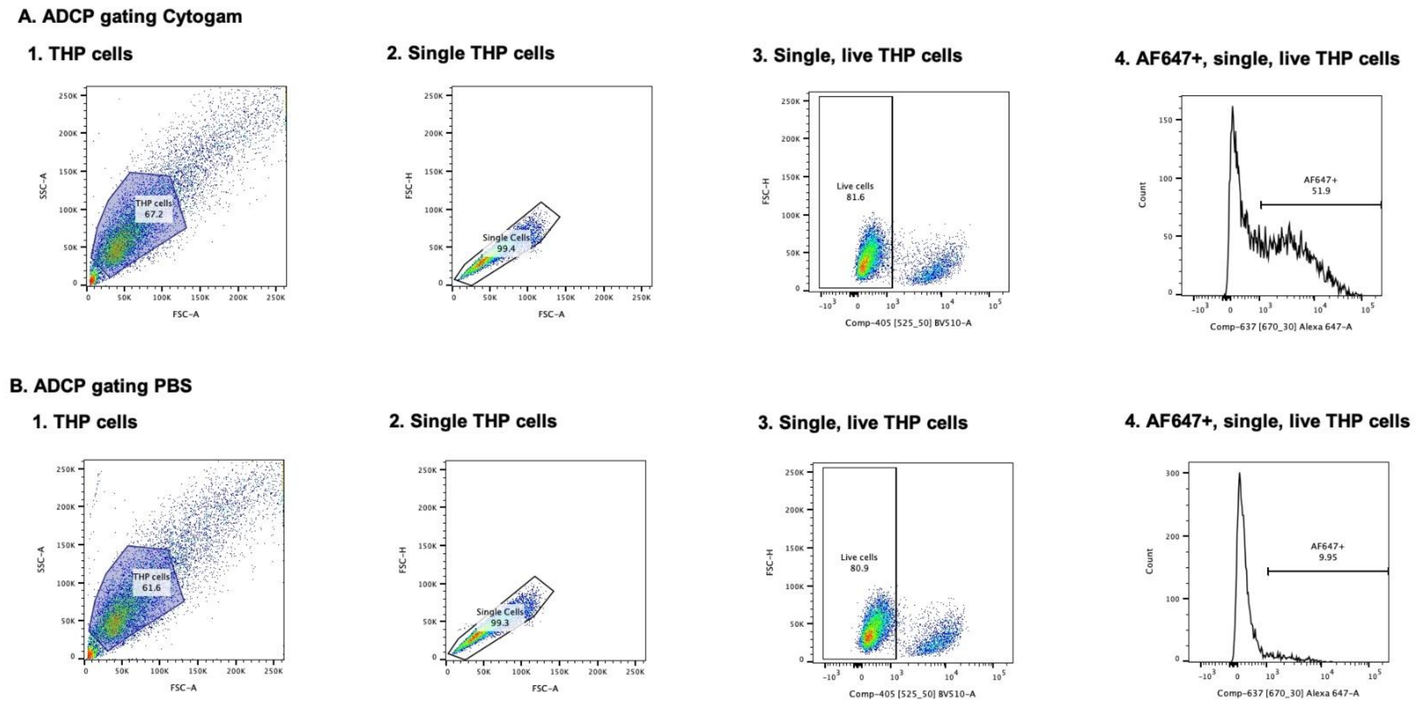


**Supplementary Figure 1: Virion phagocytosis gating strategy.** Flow cytometry gating strategy for (A) Cytogam at 50mcg/mL (positive control) and (B) PBS. FlowJo was used to gate for the percentage of single, live, AF647+ THP monocytes.
